# A salmon fish model reveals genetic signals in brain associated with early maturation

**DOI:** 10.1101/2024.03.18.585557

**Authors:** Ehsan Pashay Ahi, Jukka-Pekka Verta, Johanna Kurko, Annukka Ruokolainen, Pooja Singh, Paul Vincent Debes, Jaakko Erkinaro, Craig R. Primmer

## Abstract

Emerging evidence suggests a link between adiposity and early maturation, potentially impacting hormonal signaling pathways governing puberty timing. Fish models have proven invaluable in studying these processes, given their genetic and physiological similarities to humans. In Atlantic salmon, in addition to being linked with environmental shifts and lipid reserves, the timing of sexual maturation also has a strong genetic basis. A gene encoding a co-factor of Hippo pathway, *vgll3*, is a major determinant of maturation timing in salmon, and the same gene was found to be associated with maturation timing in humans. Intriguingly, *vgll3* also inhibits adipogenesis in mice. Recent studies in salmon revealed correlations between *vgll3* genotypes and reproductive axis gene expression, possibly modulated through the Hippo signaling pathway. The Hippo pathway is known for its role in sexual maturation and adipogenesis and responds to environmental cues such as dietary fat and temperature. In this study, we employed a custom gene expression panel in male Atlantic salmon with different *vgll3 early* (E) and *late* (L) maturation genotypes testing components of this pathway in brain at immature and mature stages. We found increased brain expression of a major Hippo pathway kinase (*lats1b*) and melanocortin receptor encoding genes (*mc4ra* and *mc4rc*) in individuals with *early* (E) maturation genotypes of *vgll3* before maturation development of testes. Moreover, we found components and interacting partners of Hippo pathway showing differential expression in brain of individuals with *early* (E) and *late* (L) *vgll3* genotypes prior to maturation. This could indicate extensive and complex roles of Hippo pathway in brain processes required for preparing for [the onset of] maturation at upstream of reproductive axis. This study elucidates molecular mechanisms underpinning early maturation, and for the first time in fish, offering insights into detecting it at molecular level in brain before visible gonadal changes occur.

## Introduction

Early maturation in humans, commonly referred to as precocious puberty, presents a complex and multifaceted challenge. This condition involves the onset of puberty, marked by physical and hormonal changes, occurring earlier than the average age. Precocious puberty can lead to a range of physical, psychological, and social issues (Hartge 2009; Day *et al*. 2017; Leka-Emiri *et al*. 2017). Early maturation has been associated with an increased risk of certain health conditions, including metabolic disorders and cardiovascular problems, later in life (Day *et al*. 2017). A growing body of evidence suggests a potential link between adiposity and early maturation in humans (Wagner *et al*. 2012). Adiposity, characterized by excess body fat, may influence the intricate interplay of hormonal signaling pathways that govern the timing of puberty (Wagner *et al*. 2012). The precise mechanisms underlying the relationship between adiposity and early maturation are complex and multifaceted, and therefore poorly understood (Leka-Emiri *et al*. 2017). Fish models have long served as valuable tools in the study of metabolic and sexual maturation processes in humans due to their remarkable physiological and genetic similarities (Zohar 2021). These models offer distinct advantages, including a conserved endocrine system that governs metabolic and reproductive processes. The genetic homology between fish and humans enables to investigate shared molecular pathways and mechanisms underlying disorders. Sexual maturation begins in the brain, predominantly with the hypothalamus releasing gonadotropin-releasing hormone (GnRH), which prompts the pituitary gland to release LH and FSH hormones (Herbison 2016). These hormones reach the gonads, spurring sex hormone production. However, pubertal timing regulation extends beyond the hypothalamus-pituitary-gonad (HPG) axis, involving other brain regions like the pineal gland, amygdala, and prefrontal cortex, interacting with environmental cues (Sisk and Foster 2004). Fish brains are less centralized than mammalian brains in terms of anatomical structures (Arendt *et al*. 2008; Cerdá-Reverter and Canosa 2009). While the complex and interconnected organization and structure of fish brains reflect the diverse evolutionary history and adaptations of fish to various aquatic environments and lifestyles, different brain regions than in their mammalian counterparts may play more or less essential role in onset of puberty (Cerdá-Reverter and Canosa 2009; Tena-Sempere *et al*. 2012; Fontaine *et al*. 2020; Somoza *et al*. 2020). For instance, fish have the "preoptic area" with a more prominent role in production of GnRH and the pineal gland in fish play a more pivotal role in reproductive behaviors and timing than in mammals (Zohar *et al*. 2010; Tena-Sempere *et al*. 2012). In salmonid fish, the pineal gland regulates seasonal reproductive changes via photoperiod (Porter *et al*. 1998; Mobley *et al*. 2021). Overall, comprehending maturation timing in fish requires more holistic brain study including more brain regions than classic studies of HPG axis. In addition, investigations of potential molecular signals in the entire brain may provide insights on whether changes in less studied parts of the brain participate in pre-maturation signals that later trigger HPG axis. This may lead to detection of early molecular signals, which could potentially stimulate the onset of puberty prior to molecular changes in HPG axis.

The timing of sexual maturation in Atlantic salmon exhibits a strong connection with seasonal shifts in the environment and body condition (House *et al*. 2023). The allocation of lipids has a pivotal role influencing salmon age at maturity, as they require a specific lipid reserve threshold to initiate maturation (Rowe *et al*. 2011). Notably, within Atlantic salmon, a single locus encompassing the gene vestigial-like family member 3 (*vgll3*) stands as the principal genetic determinant of maturation timing, associated with over 39% of the variance in age at maturity in natural populations (Barson *et al*. 2015; Czorlich *et al*. 2018). The impact of *vgll3* on age at maturity becomes evident in male salmon as early as one year old under controlled conditions (Verta *et al*. 2020; Debes *et al*. 2021; Sinclair-Waters *et al*. 2021). Intriguingly, its human ortholog (*VGLL3*) has also demonstrated an association with age at maturity (Perry *et al*. 2014; Cousminer *et al*. 2016). In addition to its role in maturation age, vgll3 serves as an inhibitor of adipogenesis in mice (Halperin *et al*. 2013). In salmon, individuals with distinct *vgll3* genotypes exhibit seasonal variations in energy storage, suggesting a noteworthy link between energy utilization and environmental changes (House *et al*. 2023; Ahi *et al*. 2024b).

Recent investigations in Atlantic salmon have demonstrated a robust correlation between *vgll3* alleles associated with *early* (E) and *late* (L) maturation and the expression patterns of a number of crucial reproductive axis genes (Ahi *et al*. 2022, 2024a). The anticipation that *vgll3* genotype effects on reproductive axis genes may be modulated through divergent Hippo signaling pathway activation has been shown by gene co-expression network analysis (Ahi *et al*. 2023). The Hippo pathway is rapidly emerging not only as a pivotal molecular signal governing sexual maturation in vertebrates (Kjærner-Semb *et al*. 2018; Sen Sharma *et al*. 2019; Kurko *et al*. 2020) but also as a regulator in the equilibrium between adipocyte proliferation and differentiation (Ahi *et al*. 2024b). Moreover, the Hippo pathway seems to play an integral role in transducing environmental cues, such as alterations in dietary fat content and temperature, into transcriptional responses (Shu *et al*. 2019; Luo *et al*. 2020). Whilst *vgll3* expression level has been negatively associated with adipocyte production (Halperin *et al*. 2013) and gonad development (Verta *et al*. 2020), within the Hippo pathway, Vgll3 operates as a significant transcription co-factor, predominantly assuming an activating role (Hori *et al*. 2020). This dynamic interplay involves competition with another major transcription co-factor of the pathway, Yap1, which primarily functions as an inhibitor of the Hippo pathway (Kurko *et al*. 2020; Hori *et al*. 2020). Notably, the activity of Yap1, a principal transcription co-factor of the pathway, is deemed indispensable in adipogenesis (Ardestani *et al*. 2018). Collectively, these findings highlight that Atlantic salmon is an excellent natural model since its distinct *vgll3* alleles provide a functional connection between maturation and adipogenesis, and offer an opportunity to study the direct molecular mechanisms interlinking sexual maturation and energy acquisition. In this study, we sought to investigate how different *vgll3* genotypes influence transcriptional activity in the brain both before and after sexual maturation. The study examined males (since the maturation occurs on average at earlier age in salmon males than females) with different maturation phenotypes (mature and immature) and genotypes (*early* vs. *late* maturing *vgll3* genotypes).

## Materials and methods

### Fish material and tissue sampling

Individuals used in the study were from the same population (Oulujoki) and cohort used in Verta et al. (2020). This provided access to individually PIT tagged individuals with known *vgll3* genotypes (see Verta et al., (2020) for more details of crossing and rearing). For this study, male individuals were sampled at the ages of 1.5 to 2 years post fertilization (Debes *et al*. 2020). Fish were euthanized by an overdose of MS222 and the entire brain (except pituitary gland) of individual males was sampled at one of three time points, each reflecting a different maturation development stage as described below:

*Immature 1*-stage individuals were collected during the late spring period (5-21 May), exhibiting an average mass of 17.5 g (with a range of 10.2-27.4 g) and an average length of 12.1 cm (ranging from 9.1-13.6 cm). These individuals displayed no discernible signs of gonad development, as indicated by a gonadosomatic index (GSI) value of 0.

*Immature 2*-stage individuals were sampled in the summer season (4-17 July), with an average mass of 33.7 g (ranging from 21.1-76.6 g) and an average length of 17.3 cm (falling within the range of 14-18.5 cm). Some of these individuals exhibited initial indications of the onset of phenotypic maturation processes, as reflected by GSI values ranging from 0.0 to 0.4.

*Mature*-stage individuals were collected during the anticipated spawning period in early autumn (1-15 October). These individuals had an average mass of 82.7 gm (with a range of 60.3-120.6 gm) and an average length of 20.8 cm (ranging from 16.4-22.7 cm). All individuals in this group displayed well-developed gonads indicative of maturity, with a GSI exceeding 3.

Subsequently, these tissue samples were rapidly frozen using liquid nitrogen and stored at a temperature of -80°C until they were processed further.

### RNA extraction

RNA was extracted from a total of 24 samples of male brain (four *vgll3*\*EE and four *vgll3*\*LL individuals for each of the three time points) using a NucleoSpin RNA kit (Macherey-Nagel GmbH & Co. KG). The collected samples were transferred to tubes containing 1.4 mm ceramic beads from Omni International, along with Buffer RA1 and DDT (350 µl of RA1 and 3.5 µl of 1M DDT). Homogenization was performed using the Bead Ruptor Elite (Omni International) at a frequency of 30 Hz for a total of 2 min (in 6 cycles of 20 s each). The subsequent RNA extraction procedures adhered to the guidelines provided by the manufacturer, with the kit also including an integrated DNase stage to eliminate any remaining genomic DNA (gDNA). At the conclusion of this process, the RNA extracted from each individual sample was eluted using 50 µl of nuclease-free water. To assess RNA quantity, measurements were taken using the NanoDrop ND-1000 instrument (Thermo Scientific, Wilmington, DE, USA), while the quality of the RNA was evaluated through the employment of the 2100 BioAnalyzer system (Agilent Technologies, Santa Clara, CA, USA). The RNA integrity number (RIN) exceeded 7 for all samples. For the hybridization step within the Nanostring panel, a total of 100 ng of the extracted RNA from each isolation was utilized.

### Nanostring nCounter mRNA expression panel

NanoString nCounter represents a multiplex nucleic acid hybridization technology, offering a reliable and consistently reproducible method for evaluating the expression of numerous RNA molecules in a single assay (Goytain and Ng 2020). This technology bears distinct advantages for ecological and evolutionary research due to its minimal RNA input requirement, which can accommodate lower-quality RNA compared to RNA-seq. Importantly, it operates without an amplification step and is capable of detecting exceptionally low levels of RNA expression. In this study, the Nanostring panel of probes was an expanded iteration of the panel used in a previous study (Kurko *et al*. 2020), which initially focused on investigating age-at-maturity-related gene expression in Atlantic salmon. This updated panel, encompassing more than 100 additional genes, particularly concentrates on an extensive collection of Hippo pathway components and other genes directly interconnected with the Hippo pathway. The gene selection process involved both literature review and the utilization of tools, such as the IPA (Ingenuity Pathway Analysis) from Qiagen, as well as other accessible web-based tools and databases (Kurko *et al*. 2020). Notably, the panel includes probes targeting genes linked to age-at-maturity in Atlantic salmon, such as *vgll3a* on chromosome 25 and *six6a* on chromosome 9, along with their corresponding paralogs *vgll3b* on chromosome 21 and *six6b* on chromosome 1. Additionally, the panel incorporates probes for other genes functionally associated with sexual maturation, including those within the hypothalamus-pituitary-gonadal (HPG) axis. Given the presence of multiple paralogs for most candidate genes due to the duplicated Atlantic salmon genome, the paralogs of each gene of interest were also integrated, having been identified through resources like SalmoBase (http://salmobase.org/) and the NCBI RefSeq databases.

Further specifics regarding gene/paralog selection and nomenclature are documented in (Kurko *et al*. 2020). The gene accession numbers, symbols, full names, and functional classifications are provided in supplementary file 1. The analysis of mRNA expression levels for these candidate genes involved the application of the Nanostring nCounter Analysis technology (NanoString Technologies, Seattle, WA, USA). Probes designed for each gene paralog, aimed at all known transcript variants, were formulated using reference sequences from the NCBI RefSeq database. However, designing paralog-specific probes proved unfeasible for some genes due to substantial sequence similarity between paralogs. Practical execution encompassed the utilization of the nCounter Custom CodeSet for probes and the nCounter Master kit (NanoString Technologies). The RNA from each sample was denatured, followed by an overnight hybridization with the probes. Subsequent to hybridization, purification and image scanning were performed the following day.

### Data analysis

Among 9 candidate reference genes in the panel, 8 genes, including *actb*, *ef1a* paralogs (*ef1aa*, *ef1ab* and *ef1ac*), *hprt1*, *prabc2* paralogs (*prabc2a* and *prabc2b*) and *rps20*, were selected for data normalization since they showed a low coefficient of variation (CV) values across the samples. The excluded reference gene was *gapdh* as it showed very high variation (CV% > 100), so even though *gapdh* is commonly used as a reference gene in many studies, it seemed to be unsuitable for data normalization in the brain of Atlantic salmon. Following this, the raw count data obtained from Nanostring nCounter mRNA expression underwent normalization through RNA content normalization factors, calculated individually for each sample using the geometric mean count values of the selected set of reference genes. After normalization, a quality control assessment was conducted, with all samples successfully passing the predefined threshold using the default criteria of nSolver Analysis Software v4.0 (NanoString Technologies; www.nanostring.com/products/nSolver). During data analysis using the software, the mean of the negative controls was subtracted, and positive control normalization was carried out by utilizing the geometric mean of all positive controls, following the manufacturer’s recommendations. To establish a baseline signal threshold, a normalized count value of 20 was set as the background signal. Consequently, among the analyzed genes, 82 genes displayed an average signal below this threshold across the samples, leading to the consideration of 255 genes for subsequent analyses. Differential expression analysis was executed using the log-linear and negative binomial model (lm.nb function) as integrated within Nanostring’s nSolver Advanced Analysis Module (nS/AAM). Inclusion of predictor covariates in the model encompassed the maturation status and genotypes, as guided by nS/AAM’s suggestions. To mitigate multiple hypothesis testing, the Benjamini-Yekutieli method (Benjamini and Yekutieli 2001) was employed within the software, with an adjusted p-value threshold of < 0.05 deemed statistically significant (supplementary file 2). Additionally, for further exploration, log-transformed expression values were utilized in calculating pairwise Pearson correlation coefficients (r) between the gene expression of each candidate gene and GSI values across the entirety of the samples.

We employed the Weighted Gene Coexpression Network Analysis (WGCNA) R-package version 1.68, within R-package version 5.2.1, to discern gene co-expression networks (GCN) as outlined by Langfelder & Horvath, 2008. Our primary focus being the comparison of alternative *vgll3* genotypes, we harnessed all samples from both maturation statuses and time-points within each genotype as biological replicates, ensuring robust statistical power for WGCNA. To ascertain sample relationships, hierarchical clustering of samples based on gene expression was performed. The construction of coexpression networks encompassed the following stages: (1) determination and quantification of gene coexpressions through Pearson correlation coefficients, (2) establishment of an adjacency matrix with a focus on scale-free topology employing the coefficients, (3) computation of the topological overlap distance matrix via the adjacency matrix, (4) hierarchical clustering of genes using the topological overlap distance with the "average" method, (5) identification of coexpressed gene modules through employment of the cutTreeDynamic function, with a minimum module size of 10 genes, (6) allocation of colors to each module and representation of module-specific expression profiles through the principal component (module eigengene), and (7) merging of highly similar modules based on module eigengene (ME) dissimilarity, utilizing a distance threshold of 0.25, to finalize the set of coexpressed gene modules. Furthermore, we implemented a conditional coexpression analysis, akin to Singh et al., 2021, wherein coexpression networks were individually constructed for each *vgll3* genotype. This approach sought to identify the preservation of *early* maturation genotype (EE) modules within the *late* maturation (LL) network and vice versa. With a softpower of 9, an adjacency matrix was established. Lastly, to assess the preservation of modules’ density and connectivity between the reference dataset (EE) and the query dataset (LL), module preservation statistics were computed using WGCNA, following the methodology outlined by Langfelder, Luo, Oldham, & Horvath, 2011. A permutation test was implemented to iteratively shuffle genes within the query network, calculating Zscores. The individual Z scores from all 200 permutations were summarized into a Zsummary statistic.

We employed Manteia (Tassy and Pourquié 2014) for Gene Ontology (GO) enrichment analysis to identify enriched biological processes within each gene co-expression module. The GO enrichment criteria were set at FDR < 0.05, with a specific GO level 2 threshold for inclusion. For the anticipation of potential gene interactions and the identification of key genes displaying the highest interaction count (referred to as interacting hubs), the differentially expressed genes identified in each comparison were converted to their conserved orthologs in humans. These orthologs were chosen due to their extensive validated and studied interactome data across vertebrates. Subsequently, these orthologs were used as input for STRING version 12.0, a comprehensive knowledge-based interactome database for vertebrates (Szklarczyk *et al*. 2023). Predicted gene interactions were based on multiple factors such as structural similarities, cellular co-localization, biochemical interactions, and co-regulation. The confidence level for each interaction or molecular connection prediction was maintained at a medium setting, which is the default threshold.

## Results

### Differences in gene expression between *vgll3* genotypes

We first evaluated variation in expression between different *vgll3* genotypes during distinct developmental stages: *Immature1* and *Immature2* (late spring and summer periods), as well as *Mature* (early autumn) (Fig. 1). We found 8 genes expressed differentially between genotypes at the *Immature 1*, and 13 genes at the *Immature 2* time point, as well as 19 genes at the *Mature* time point. At *Immature 1*, all differentially expressed genes, except *lats1b* and *mc4rc*, exhibited lower expression in *vgll3*EE* genotype individuals (Fig. 1A). The query of molecular interactions revealed that while none of the genes had direct interactions with *vgll3*, three genes showed potential direct interaction with *yap1* (*amotl1, lats1b* and *snai1*) (specified with connecting lines between the genes in Fig. 1B). At *Immature 2*, we found 10 genes with higher expression in *vgll3*EE* genotype individuals and 3 genes showing higher expression in *vgll3*LL* genotype individuals (Fig. 1C). The predicted interactions revealed 4 genes (*egr1a*, *foxo1c*, *snai2b* and *sox9c*) with direct molecular interaction with *yap1* whereas again no gene had direct interaction with *vgll3* (Fig. 1D). Furthermore, all of the 4 genes showing interaction with *yap1* also formed interacting hubs with other differentially expressed genes (based on them showing the highest number of interactions compared to other genes) and *egr1a* displayed the highest number of connections indicating its potential key functional role in this interaction network (Fig. 1D). Moreover, *erg1a* was the only gene among the 4 genes interacting with *yap1* showing lower expression in *vgll3*EE.* Finally, at *Mature*, we found 5 genes with higher expression in *vgll3*EE* genotype individuals, and 14 genes showed higher expression in *vgll3*LL* genotype individuals (Fig. 1E). The interaction query identified 6 genes (*egr1d*, *kdm5b*, *lats2a*, *rhoad*, *rnd3b*, and *wnt5a*) with direct interactions with *yap1*, whereas one gene (*kdm5b*) had a direct interaction with *vgll3* (Fig. 1F). Except for *lats2a*, the remaining genes showing direct interactions with *yap1* had lower expression in *vgll3*EE* individuals and one of them, *rhoad*, formed an interacting hub, i.e. had highest number of interactions with other differentially expressed genes (Fig. 1F).

**Figure 1:**
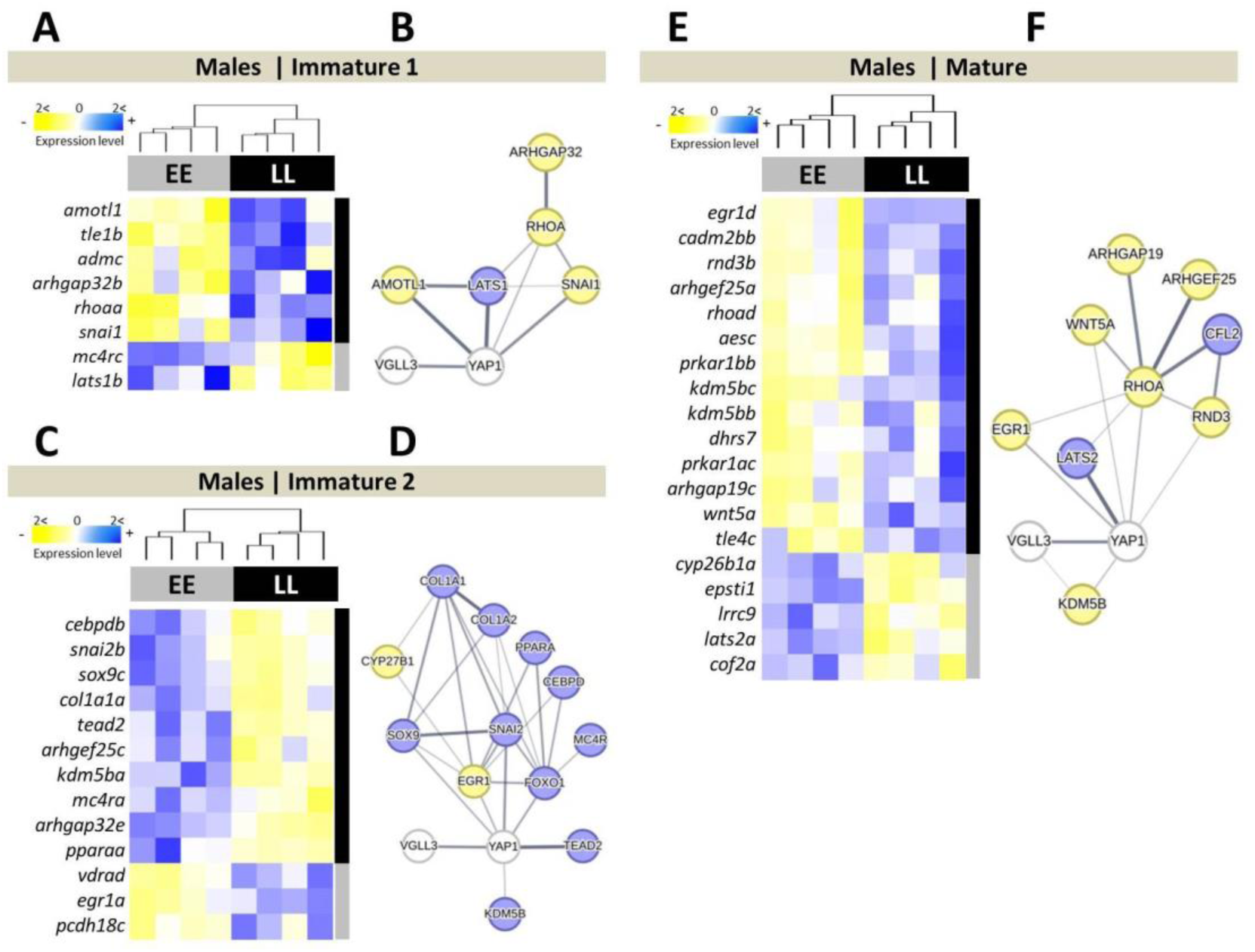
Differentially expressed genes between alternative *vgll3* genotypes and their predicted interactions in Atlantic salmon brain. Heatmaps represent differentially expressed genes between *vgll3* genotypes at three timepoints (A, C, E) and their respective predicted molecular interactions (B, D, F). The thickness of the connecting lines between the genes indicates the probability of interaction. Genes colored in blue and yellow in the predicted network indicate higher and lower expression in *vgll3*EE* individuals, respectively.

### Maturation specific gene expression differences

To pinpoint alterations in gene expression linked to the transition from an immature to a mature state, we compared immature individuals obtained at both *Immature 1* and *2* time points and those collected at the *Mature* time point. We found 20 differentially expressed (DE) genes between mature and immature individuals both when *vgll3* genotypes were combined, and when *vgll3* genotypes were analyzed separately, 8 and 25 DE genes in *vgll3*LL* and *vgll3*EE* genotypes, respectively (Fig. 2A-C). In *vgll3*LL* individuals, 6 out of the 8 and in *vgll3*EE* individuals 11 out of the 25 DE genes between immature vs mature individuals had higher expression at the *Mature* time point (Fig. 2B-C). Across all three comparisons, even though 12 genes showed differential expression when both genotypes were combined, no gene was found to be differentially expressed between the stages independent of *vgll3* genotype (Fig. 2A-D). We proceeded to delve deeper into potential functional/molecular interactions among differentially expressed genes that exhibited *vgll3* genotype-specific variation in expression (colored numbers in Fig. 2D). The predicted interactions between these genes revealed that in the *vgll3* genotype specific comparisons, 4 and 14 DE genes within *vgll3*LL* and *vgll3*EE* individuals, respectively, were found to be part of a common interaction network (colored green and red in Fig. 2E). Among the DE genes in *vgll3*LL* individuals, 3 genes, *kdm5ba*, *lats1b* and *stk3b*, had direct interactions with *yap1* and all showed higher expression in the mature stage within this genotype (Fig. 2E). Among the DE genes in *vgll3*EE* individuals 5 genes, *cdh2d*, *egr1c/d*, *tead1* and *tead2*, had direct interactions with *yap1* and also two of them, *tead1* and *tead2*, had direct interactions with *vgll3*. Other genes with a high number of interactions in the network include *ajubaa*, *rnd3d*, *rxrab* and *pitx1a*, all of which were differentially expressed in *vgll3*EE* and all except *pitx1a* had higher expression in the *mature* stage) (Fig. 2E).

**Figure 2:**
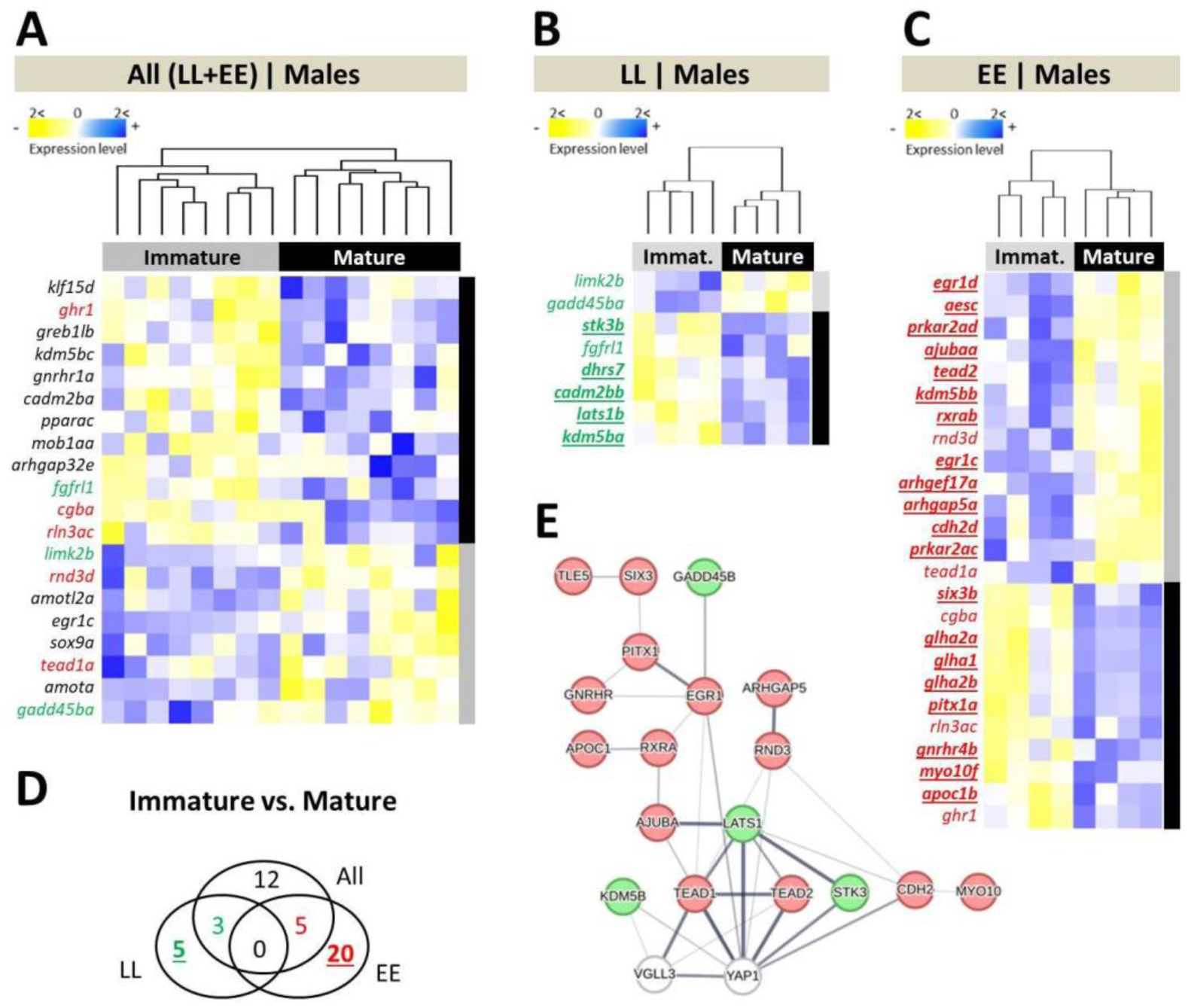
Differentially expressed genes between immature versus mature male Atlantic salmon and their predicted interactions in brain. Heatmaps representing differentially expressed genes between the immature versus mature males across alternative *vgll3* homozygotes (**A**), and within *vgll3*LL* (**B**) and *vgll3*EE* genotype individuals (**C**). A Venn diagram showing the numbers of differentially expressed genes overlapping between the comparisons (**D**). Predicted interactions between the genes colored in green and red indicating the overlapping genes in the Venn diagram (**E**). The thickness of the connecting lines between the genes indicates the probability of the interaction.

Subsequently, we examined the expression of each gene of the panel in correlation with the gonadal development stage (GSI), aiming to uncover genes exhibiting expression profiles within the brain that are closely intertwined directly with the gonadal maturation process. We observed a positive correlation between the expression of three genes—*ghr1*, *klf15d*, and *cebpba*—and GSI, regardless of the *vgll3* genotype. (Fig. 3A-D). Furthermore, two of these genes, *ghr1* and *klf15d*, were among the genes with most significant positive expression correlations within each genotype. Overall, the majority of the correlations detected with GSI were positive, irrespective of the grouping of *vgll3* genotypes (Fig. 3A-C). Four genes amongst *vgll3*EE* genotype individuals, *arhgap19a/b*, *myo10e* and *snaib*, showed negative expression correlations with GSI, and all were shared between the comparisons of *vgll3*EE* genotype and both genotypes together. Amongst *vgll3*LL* individuals, expression of one gene, *ets1c*, was found to be negatively correlated with GSI (Fig. 3B). The predicted interactions between the genes showing genotype-specific expression correlations (colored numbers in Fig. 3D) revealed a potential network between 2 and 5 genes, respectively, for *vgll3*LL* and *vgll3*EE* genotype individuals, as well as 2 overlapping genes across all groups (Fig. 3E). For *vgll3*EE* genotype individuals, 2 genes (*snaib* and *mob1ab*) showed direct interactions with *yap1* and one gene (*greb1lb*) with *vgll3,* whereas for *vgll3*LL* genotype only one gene, *ets1c*, showed a direct interaction with *vgll3*.

**Figure 3:**
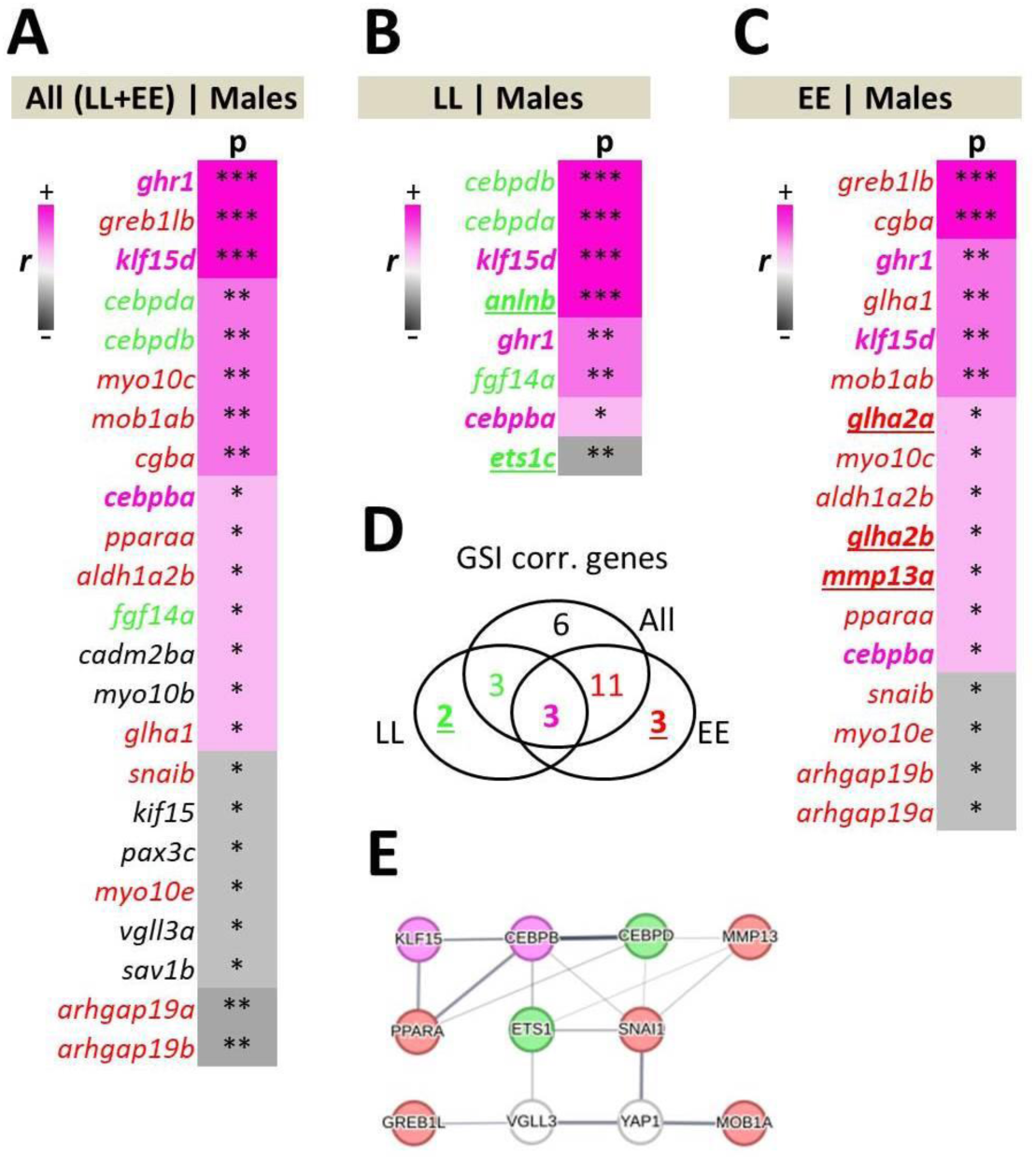
Genes showing GSI correlated expression and their predicted interactions in brain. Ranking of significant Pearson correlations between gene expression and GSI in salmon brain across both *vgll3* genotypes (**A**), within *vgll3*LL* (**B**) and *vgll3*EE* (**C**) genotype individuals. The Venn diagram depicts the numbers of genes significantly correlated with GSI unique or overlapping between the comparisons (**D**). Predicted interactions between the genes colored in green, red and purple indicate the overlapping genes in the Venn diagram (**E**). EE and LL indicate *vgll3*EE* and *vgll3*LL* genotypes, respectively, and p and r imply on p-values (* < 0.05; ** < 0.01; *** < 0.001) and Pearson correlation coefficient. The thickness of the connecting lines between the genes indicates the probability of the interaction.

### Identification of gene coexpression networks

To attain a more comprehensive understanding of the transcriptional dynamics involving components of the Hippo pathway and their established interacting genes, we employed network-based co-expression analyses. This approach facilitated the monitoring of genotype-specific alterations within each network. To execute this strategy, we initially constructed gene coexpression networks (GCN) in the brain of each *vgll3* genotype. Subsequently, we evaluated the extent to which the identified gene co-expression modules were conserved between the genotypes. In essence, we established the GCN within one genotype (*vgll3*EE* or *vgll3*LL*) and subsequently tested the preservation of its modules within the other genotype (*vgll3*LL* or *vgll3*EE*), respectively.

We identified 4 modules within *vgll3*EE* GCN (Fig. 4A and B) of which 2 modules, brown and yellow, showed low preservation (Zsummary < 2) in *vgll3*LL*, *i.e.* some of the genes in each module do not have significant expression correlations in *vgll3*LL* genotype (Fig. 4A). For each module, we performed an enrichment analysis of Gene Ontology (GO) Biological Processes to unveil the major processes in which the genes within each module are engaged. In the yellow module, we found associations with processes of regulation of hormone levels, signal transduction and fat-soluble vitamin metabolism (Fig. 4C). The yellow module showed the least preservation between the genotypes with 11 out of its 23 genes showing no coexpression preservation in *vgll3*LL* genotype individuals (genes lacking colour in each module in Fig. 4C). In the brown module, we found associations with processes of positive regulation of transcription and the Hippo pathway signaling and only 5 out of 25 genes showing no coexpression preservation in *vgll3*LL* genotype individuals (Fig. 4C). The removal of unpreserved genes in the yellow and brown modules led to loss of significance of two GOs in yellow and one GO in the brown module; regulation of hormone levels and signal transduction in the yellow and Hippo signaling in the brown modules (non-colored GOs in Fig. 4C).

**Figure 4.**
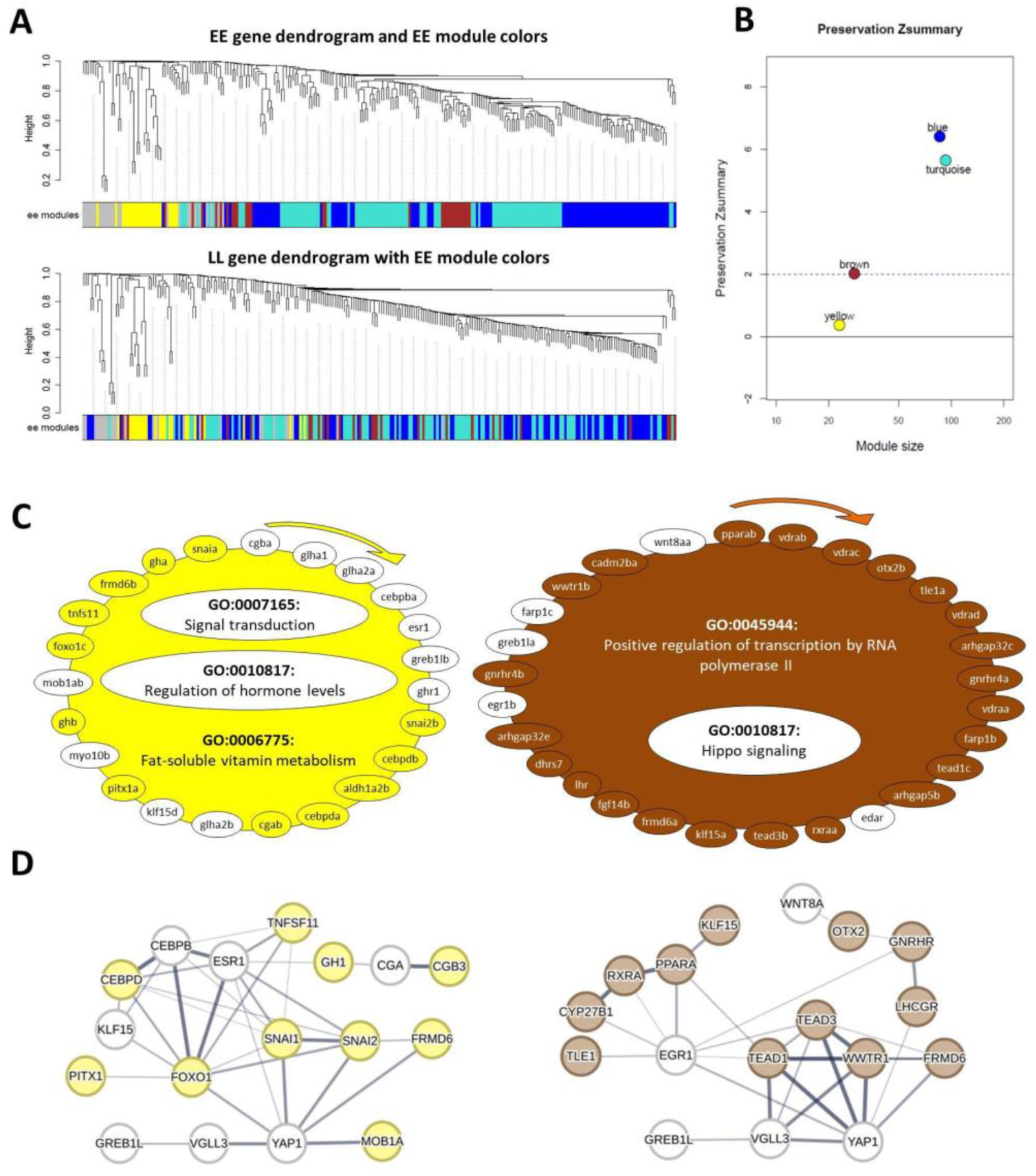
Coexpression analysis of *vgll3*EE* genotype in brain of male Atlantic salmon. (A) Visual representation of *vgll3*EE* module preservation in *vgll3*LL* individuals. The dendrograms represent average linkage clustering tree based on topological overlap distance in gene expression profiles. The lower panels of the dendrograms represent colors that correspond to the *vgll3*EE* clustered coexpression modules. Top: *vgll3*EE* modules with assigned colors. Bottom: visual representation of the lack of preservation of *vgll3*EE* module genes in *vgll3*LL* genotype individuals. (B) Preservation Zsummary scores in the *vgll3*LL* for *vgll3*EE* modules (colours represent *vgll3*EE* modules). Zsummary < 2 represents lack of preservation (dotted blue line) and Zsummary between 2 and 10 implies moderate preservation. (C) The genes in each of the two identified modules for *vgll3*EE* genotype with least preservation in *vgll3*LL*. The genes without color in each module are those showing no preserved expression correlation in *vgll3*LL* individuals and the clockwise arrows above each module indicate the direction of genes with highest to lowest expression correlations with other genes within that module. For each of the modules, the top enriched GOs are represented, and GOs without color were no longer enriched after removal of the genes without colors. (D) Predicted interactions between the genes within each of the less preserved modules. Different levels of thickness in the connecting lines between the genes indicates the probability of the interaction.

Utilizing knowledge-based interactome prediction analysis, we employed the genes contained within each module to unveil potential interactions among these genes, along with the identification of hub genes exhibiting the greatest number of interactions. The prediction of interactions between the genes within each low preserved module revealed that in both modules at least one gene among those not showing coexpression preservation had direct interaction with *vgll3* (represented with lines directly connecting the non-colored genes with *vgll3*) (Fig. 4D). Interestingly, in each module a different paralog of *greb1l* (*greb1lb* in the yellow and *greb1la* in the brown modules) was in direct interaction with *vgll3* but lacking coexpression preservation in *vgll3*LL* genotype individuals (Fig. 4D). Moreover, in each module one gene with direct interaction with *yap1* also lacked coexpression preservation in *vgll3*LL* genotype individuals (*esr1* in the yellow and *egr1b* in the brown modules) (Fig. 4D).

In *vgll3*LL* individuals, we found only 2 modules and one of them (the blue module) showed a low level of preservation (Zsummary < 2) compared to *vgll3*EE* individuals (Fig. 5A and B). The other module (turquoise module) was very large with 215 co-expressed genes and also with the highest preservation level between the genotypes (Zsummary > 10). The blue module included genes involved in positive regulation of transcription and regulation of hormone levels (Fig. 5C), indicating potential interactions between these processes in *vgll3*LL* individuals. In *vgll3*EE* individuals, however, these two processes were already dissociated in different modules (Fig. 4C), and similarly, we observed that such a connection disappeared after the removal of genes showing no expression correlations in *vgll3*EE* (genes without color in Fig. 5C). This may also indicate more extensive connections between the general increase of transcriptional activity and hormone production in *vgll3*LL*, whereas increase of transcriptional activity in *vgll3*EE* seemed to be linked to activity of Hippo pathway (Fig. 4C and 5C). In other words, the activity of Hippo pathway might be less affected and/or might play a less extensive role in the brain of *vgll3*LL* individuals compared to *vgll3*EE* individuals. The prediction of interactions between the genes within the blue module revealed that 3 genes have direct interactions with *vgll3* and *yap1*, however, only one of the genes interacting with *yap1* (*egr1b*) has lost its coexpression preservation in *vgll3*EE* (Fig. 5D).

**Figure 5.**
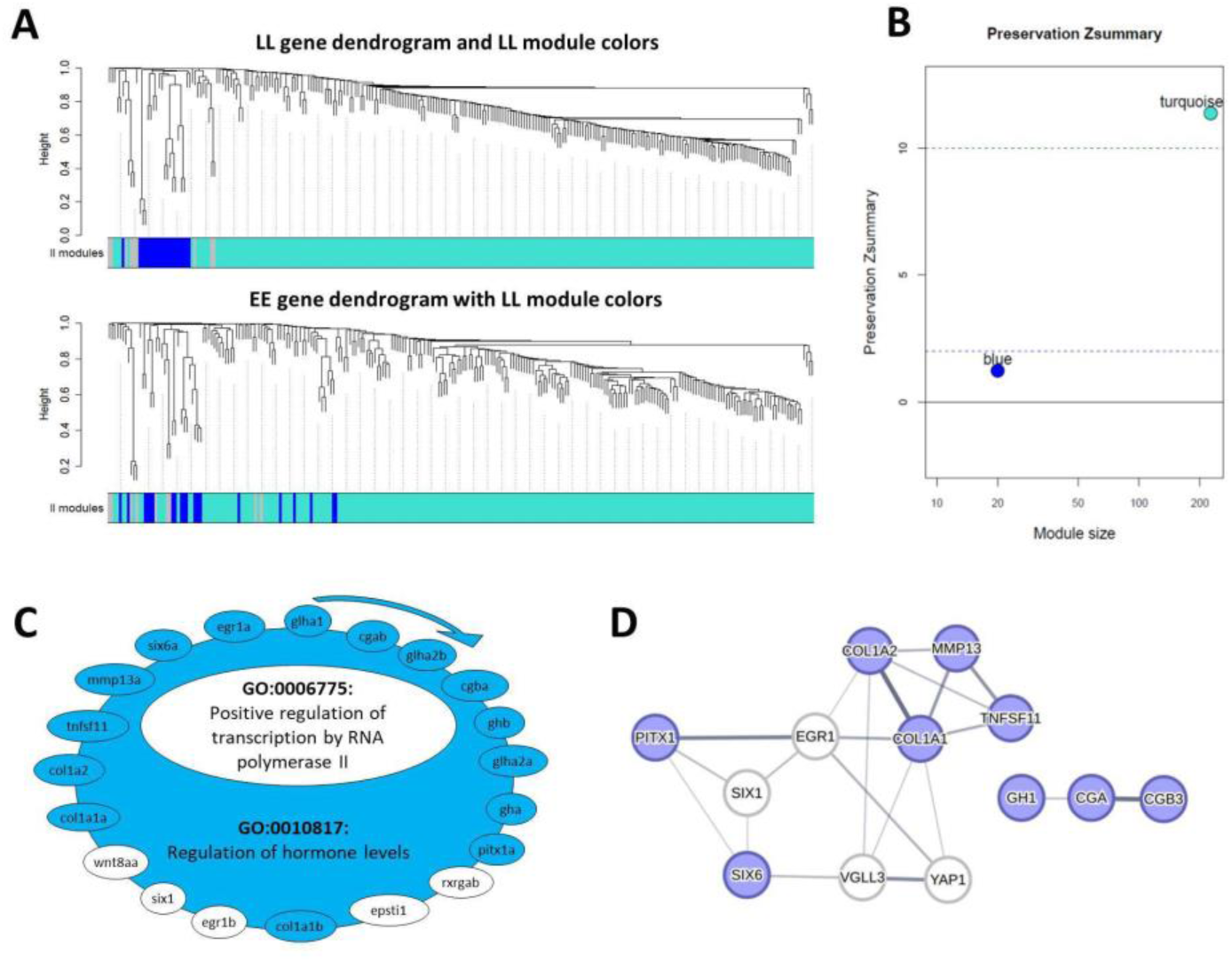
Coexpression analysis of *vgll3*LL* genotype in brain of male Atlantic salmon. (A) Visual representation of *vgll3*LL* module preservation in *vgll3*EE* individuals. The dendrograms represent average linkage clustering tree based on topological overlap distance in gene expression profiles. The lower panels of the dendrograms represent colors that correspond to the *vgll3*LL* clustered coexpression modules. Top: *vgll3*LL* modules with assigned colors. Bottom: visual representation of the lack of preservation of *vgll3*LL* module genes in *vgll3*EE* genotype individuals. (B) Preservation Zsummary scores in the *vgll3*EE* for *vgll3*LL* modules (colours represent *vgll3*LL* modules). Zsummary < 2 represents lack of preservation (dotted blue line) and Zsummary between 2 and 10 implies moderate preservation. (C) The genes in the identified module for *vgll3*LL* genotype with least preservation in *vgll3*EE*. The genes without color in the module are those showing no preserved expression correlation in *vgll3*EE* individuals and the clockwise arrows above the module indicate the direction of genes with highest to lowest expression correlations with other genes within the module. Within the module, the top enriched GOs are represented, and the GO without color was no longer enriched after removal of the genes without colors. (D) Predicted interactions between the genes within the less preserved module. Different levels of thickness in the connecting lines between the genes indicates the probability of the interaction.

## Discussion

In this study, our aim was to uncover how different genotypes of *vgll3* gene, a key determinant of sexual maturity age in Atlantic salmon (Barson *et al*. 2015), as well as a critical co-factor of the Hippo pathway (Hori *et al*. 2020), influence the transcriptional activity in the brain prior and after sexual maturation. Furthermore, we explored whether these *vgll3*-associated transcriptional changes in the brain could have the capacity for detecting early maturation before any visible signs of maturation emerge in the testis. This is particularly interesting because the molecular signal regulated by *vgll3* (the Hippo pathway) is active and plays different roles in all the brain regions involved in sexual maturation (Lodge *et al*. 2016; Beck and Kressel 2020; Lalonde-Larue *et al*. 2022). In addition, the Hippo pathway is the mediator of fatty diet-induced transcriptional changes in different tissues including brain and particularly hypothalamus (Poon *et al*. 2013). Here, we studied the expression variation of genes encoding components of the Hippo pathway and their associated interacting partners/signals, along with genes involved in sexual maturation, in the brains of individuals with different *vgll3* maturation alleles (early or late) using a specially designed Nanostring platform. This approach enabled us to anticipate how Hippo pathway signaling is regulated in the brain of Atlantic salmon, a species which exhibits a breeding strategy closely linked to seasonal changes and adiposity. Also, we could explore if components of a pivotal pathway with various roles in the brain can be detected during early maturation. Figure 6 presents a synthesis of the most important findings of this study.

**Figure 6.**
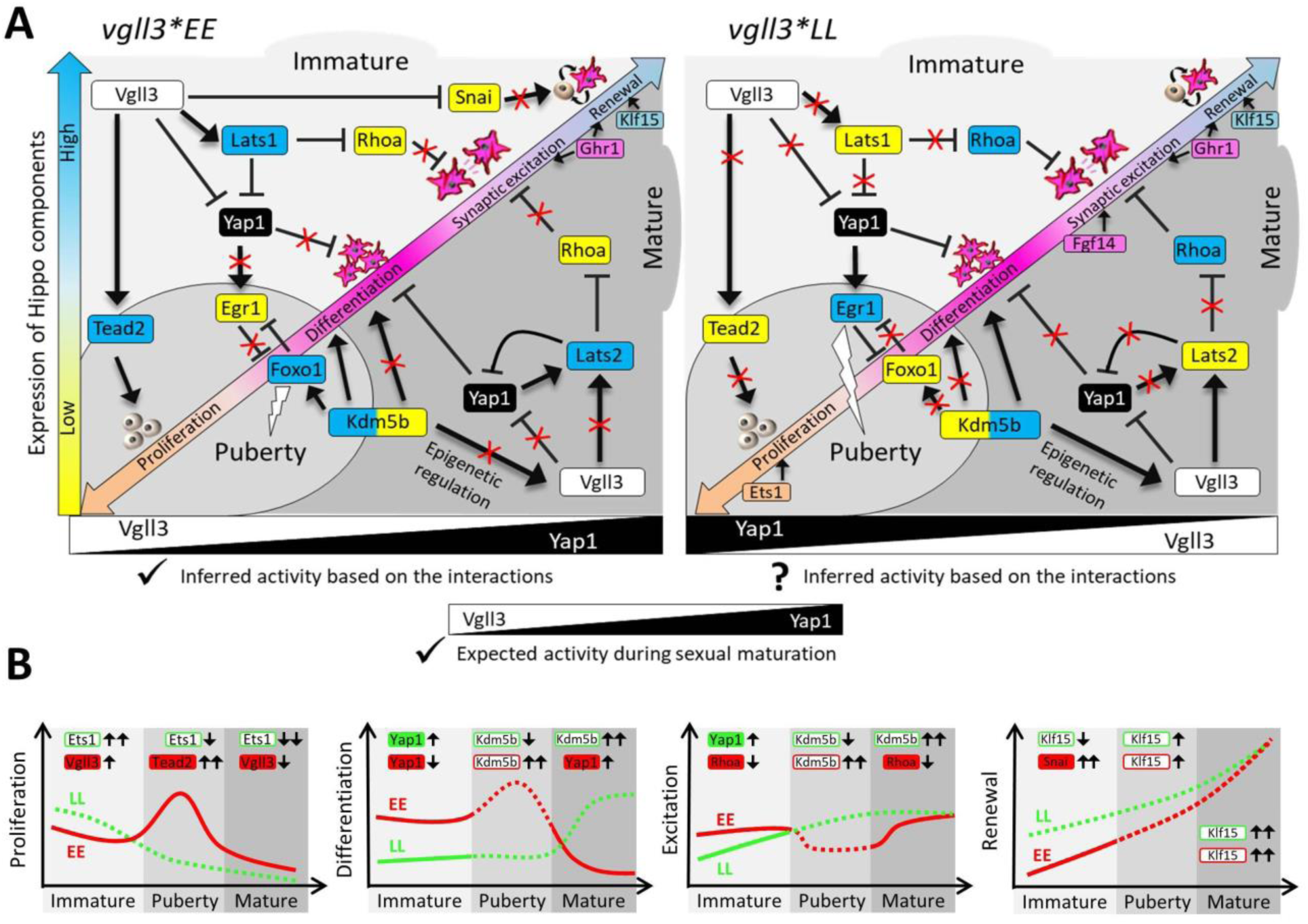
A schematic representation of the main findings linking *vgll3* genotype and Hippo pathway components to the central onset of sexual maturation in the Atlantic salmon brain. (A) Predicted gene regulatory interactions of Hippo pathway components and their interacting partners in the brain of Atlantic salmon with distinct *vgll3* genotypes. Yellow and blue colors indicate higher and lower expression of genes within the Hippo signaling pathway, respectively. The trans-sectional colored texts indicates different functional aspects of neural cells during sexual maturation and genes with related color codes to each function indicating their expression correlation with different maturation statuses. The arrow and blocked heads represent positive and negative regulations, respectively, and red crosses indicate absence of the regulatory connections. (B) Summary graphs representing each of the functions in neural cells during the sexual maturation based on the differential expression of related gene markers in each vgll3 genotypes. Genes are color coded according to *vgll3* genotype (green or red), and filled genes are major components of Hippo pathways whereas white filled genes are interacting partners of the pathway. The solid and dashed lines indicate Hippo pathway dependency and independency of the changes, respectively.

### Detection of *vgll3*-dependent maturation signals in the brain prior to the onset of puberty

We found *lats1b*, a paralog of a gene encoding a major kinase in Hippo pathway to have higher expression in the brain of individuals with the *vgll3*EE* genotype in the earliest immature stage samples (*Immature-1*). At the core of the Hippo pathway lies a series of kinase reactions (Kurko *et al*. 2020). Activation of major kinases (Lats1/2) in this pathway phosphorylates and deactivates Yap1 and thus prevents the movement of Yap1 into the nucleus, thereby governing critical genes associated with cell division, apoptosis, and cell migration (Zhang *et al*. 2008). Although this molecular process is highly conserved in animals (Zhang *et al*. 2008), it is not known if the suppression of Yap1 activity by Lats1 plays a role in early signals of triggering sexual maturation in brain. In rat, high fat diet decreases the level of Yap1 activity and subsequently increases the neural differentiation in the hypothalamus (Poon *et al*. 2013). This may indicate that individuals with the *vgll3*EE* genotype have an overall higher level of neural differentiation due to lats1b mediated inactivation of yap1 in the brain (Figure 6). This is extremely noteworthy as it suggests that the onset of early maturation is detectable in the brain prior to morphological changes in the gonads. This is because hypothalamic neural differentiation is an essential cellular process prior to puberty, which leads to fully active HPG axis and the downstream neuroendocrine cascade ensuring gonadal activation (Naulé *et al*. 2021).

Another striking finding was higher expression of *mc4rc*, a paralog of gene encoding a membrane-bound receptor and member of the melanocortin receptor family, in the brain of individuals with *vgll3*EE* genotype again at *Immature-1* (similar to *lats1b*). In certain fish species, such as *Xiphophorus* swordtails and medaka, the Melanocortin 4 receptor (*Mc4r*) has been linked to the initiation of puberty (Liu *et al*. 2019, 2020). *Mc4r* belongs to the class A of G-protein coupled receptors (GPCR) and plays a role in maintaining energy balance. Notably, in humans, mutations in the *MC4R* gene represent the most prevalent monogenic cause of severe early-onset obesity (Farooqi *et al*. 2003). Thus, increased expression of *mc4rc* in the brain of immature males could be another interesting candidate for detection of early maturation long before changes in HPG axis, however, the potential molecular link between *vgll3-*mediated Hippo pathway activation and *mc4r* has not been investigated in any species. Finally, we also found reduced expression of two major factors *snai1* and *rhoaa* in *vgll3*EE* genotype individuals. *Snai1* is an activator of yap1 and thus it can lead to similar results as above, i.e., early onset of maturation, but independent of lats1 (Tang *et al*. 2016). The function and expression of *rhoaa* is suggested to be inhibited by *vgll3*-dependent activation of Hippo pathway (Kurko et al., 2020). Also in humans, mutations in Rhoa have been recently suggested to be linked with central precocious puberty (Sklirou and Lahr 2023), and hence, it can be another conserved molecular player indicating early maturation in the vertebrate brain (Figure 6).

### Distinct interacting partners of the Hippo pathway may participate in the onset of puberty in the brain

In the later immature time-point, *Immature-2*, close to the onset of puberty (from no signs to initial signs of gonadal changes), we found a major component and several interacting partners of the Hippo pathway to be differentially expressed. A major transcription factor of this pathway, *tead2*, had higher expression in the brain of individuals with *vgll3*EE* genotype. *Tead2* is one of the highly expressed components of the Hippo pathway in mouse brain and its expression is associated with neural cell proliferation (Sahu and Mondal 2021). One of the interesting interacting partners of Hippo pathway with higher expression in the brain of *vgll3*EE* individuals was a gene encoding a paralog of Forkhead box protein O1 (*foxo1c*). In mouse, *Foxo1* encodes a transcription factor that participates in regulation of the onset of puberty by affecting the release of GnRH in the hypothalamus (Liu *et al*. 2022), however, such a role has to our knowledge not yet been reported in fish species. Two other interacting partners of the Hippo pathway with potential roles in central regulation of the onset of puberty were *kdm5b (jarid1b)*, which has been shown to be modified epigenetically prior to puberty in the rat brain (Lomniczi *et al*. 2015), and *mc4ra*, another paralog of Melanocortin 4 receptor (*Mc4r*) with conserved role in stimulating the central initiation of the puberty (see above). The only interacting partner of Hippo with lower expression in *vgll3*EE* individuals was a paralog encoding an immediate early gene (IEG) called early growth factor-1 (*egr1a*). Strikingly, *egr1* is known to be a major factor mediating socially-induced sexual maturation in fish hypothalamus (Maruska and Fernald 2011), and also we have recently found its differential expression as part of a pituitary gene network associated with *vgll3*-depenendent transcriptional regulation of gonadotropins (Ahi *et al*. 2023). However, since *egr1* is a positive regulator of sexual maturation in the HPG axis, its higher expression in *vgll3*LL* individuals may indicate a different path for those harboring *late* alleles in sexual maturation which may more relate to social interactions. A recent study demonstrated that Atlantic salmon with the *vgll3*LL* genotype show higher levels of aggression (Bangura *et al*. 2022), and interestingly, the socially induced sexual maturation through *egr1* is linked to social interactions involving aggressive/submissive (rather than cooperative) behaviors in cichlid fish (Kasper *et al*. 2018). Taken together, these findings may suggest different interacting partners of Hippo pathway participating in the onset of *vgll3*-dependent sexual maturation in Atlantic salmon (Figure 6).

### Maturation progression involves different components of the Hippo pathway in alternative *vgll3* genotypes

The direct expression comparison between immature and mature stages suggests distinct components of the Hippo pathway are involved in the maturation process in the alternative *vgll3* homozygotes. In *vgll3*LL* individuals during the transition, two genes encoding major components of the Hippo pathway, *lats1b* and *stk3b*, are predicted to have direct interactions with *yap1* and also showed higher expression in the mature than the immature stage. Lats1b is a critical regulator of Yap1 activity as described above (Zhang *et al*. 2008). While its expression was higher in *vgll3*EE* individuals at the immature stage, its role seems to be important during maturation in *vgll3*LL* individuals indicating a time-dependent expression shift (heterochrony) for *lats1b* in the brain of individuals with distinct genotypes, and probably resulting heterochrony in the neural differentiation (Poon *et al*. 2013). Interestingly, *stk3a* encodes a highly conserved activator of Lats1b (so called Mst2), which is essential for Lats1b mediated inhibition of Yap1 (Piccolo *et al*. 2014), and therefore provide stronger evidence for presence of Mst2-Last1b-Yap1 regulatory axis in the brain of *vgll3*LL* individuals upon entrance to maturation state. During the transition to maturation in the brain of *vgll3*EE* individuals, two other major regulators of Hippo pathway, *tead1* and *tead2*, which also have direct interaction with both *vgll3* and *yap1*, had reduced expression suggesting lower level of neural cell proliferation (but not differentiation) in the mature stage (Sahu and Mondal 2021).

## Conclusions

This study provides important molecular evidence linking brain gene expression and genotype-linked life-history strategy variation (sexual maturation). We thereby present gene expression evidence supporting the detection capacity of early maturation at the central nervous system level prior to the emergence of gonadal changes in Atlantic salmon immature males. This is represented by distinct transcriptional signature of components/interacting partners of the Hippo pathway, which seems to be significantly affected in individuals with early and late genotypes of *vgll3* - a main co-factor of this pathway. Furthermore, such differences seem to maintain until maturation stage, however, the involved Hippo pathway components appeared to change within and between the genotypes and maturation stages. These findings suggest that specific elements within the Hippo signaling pathway could play a significant role in causing the varying impacts of *vgll3* genotypes on brain gene expression and the timing of maturation. Nevertheless, more comprehensive functional evaluations are necessary to further confirm these proposed distinctions.

## Data availability

The authors affirm that all data necessary for confirming the conclusions of the article are present within the article, figures, and tables.

## Acknowledgements

We thank N. Piavchenko, S. Andrew, O. Andersson, T. Aykanat, Y. Czorlich, A. House, M. Lindqvist, N. Lorenzen, O. Mehtälä, K. Mobley, J. Moustakas-Verho, O. Ovaskainen, S. Papakostas, N. Parre, V. Pritchard, K. Salminen, M. Sinclair-Waters, S. Tillanen, and K. Zueva for help related to gamete stripping, sample processing, tagging, genotyping, phenotyping or fish husbandry, and the staff at the Natural Resources Institute Finland (Luke) hatchery in Taivalkoski hatchery for help during spawning.

## Author Contributions

EPA, CRP, JPV, JK and PD conceived the study; JPV, PD and CRP reared and sampled the fish; CRP and JE provided resources; AR, EPA and JPV performed experiments; EPA, JPV, PS and JK developed methodology and analyzed the data; EPA, JPV and CRP interpreted results of the experiments; EPA, JPV and CRP drafted the manuscript, with EPA having the main contribution, and all authors approved the final version of manuscript.

## Ethical approval

Animal experimentation followed European Union Directive 2010/63/EU under license ESAVI/35841/2020 granted by the Animal Experiment Board in Finland (ELLA).

---

**Supplementary File 1: Information about the gene probes on the NanoString panel.**

**Supplementary File 2: Expression data and statistical analysis.**

